# Kinematic markers of skill in first-person shooter video games

**DOI:** 10.1101/2023.02.27.530169

**Authors:** Matthew Warburton, Carlo Campagnoli, Mark Mon-Williams, Faisal Mushtaq, J. Ryan Morehead

## Abstract

Video games present a unique opportunity to study motor skill. First-person shooter (FPS) games have particular utility because they require visually-guided hand movements that are similar to widely studied planar reaching tasks. However, there is a need to ensure the tasks are equivalent if FPS games are to yield their potential as a powerful scientific tool for investigating sensorimotor control. Specifically, research is needed to ensure that differences in visual feedback of a movement do not affect motor learning between the two contexts. In traditional tasks, a movement will translate a cursor across a static background, whereas FPS games use movements to pan and tilt the view of the environment. To this end, we designed an online experiment where participants used their mouse or trackpad to shoot targets in both contexts. Kinematic analysis showed player movements were nearly identical between conditions, with highly correlated spatial and temporal metrics. This similarity suggests a shared internal model based on comparing predicted and observed displacement vectors, rather than primary sensory feedback. A second experiment, modelled on FPS-style aim-trainer games, found movements exhibited classic invariant features described within the sensorimotor literature. We found that two measures of mouse control, the mean and variability in distance of the primary sub-movement, were key predictors of overall task success. More broadly, these results show that FPS games offer a novel, engaging, and compelling environment to study sensorimotor skill, providing the same precise kinematic metrics as traditional planar reaching tasks.

**Significance statement:** Sensorimotor control underpins human behaviour and is a predictor of education, health, and socioemotional wellbeing. First-person shooter (FPS) games hold promise for studying sensorimotor control at scale, but the visual feedback provided differs from traditional laboratory tasks. There is a need to ensure they provide measures that relate to traditional tasks. We designed an experiment where the visual contingency of movements could be varied whilst participants shot targets. Participant’s movements were similar between contexts, suggesting the use of a common internal model despite the sensory differences. A second experiment observed canonical learning patterns with practice and found two measures of mouse control strongly predicted overall performance. Our results highlight the opportunity offered by FPS games to study situated skilled behaviour.

## Introduction

Video games are an increasingly ubiquitous form of entertainment, with many gamers so skilled they are employed to play in tournaments where hundreds of millions of dollars are awarded every year (1). Action video games (encompassing first-person shooter [FPS] games) are especially popular, and typically engage a range of cognitive, perceptual, and motor abilities (2, 3). The links between gaming and perceptual and cognitive abilities have been well studied, finding that action video game players are better attuned to perceptual variables (4, 5), have lower reaction times (6, 7), and a greater ability to direct visual attention (8, 9), among many other faculties (2, 10). Few studies, however, have focussed on skill development in these games, and in particular the role of the motor system in reaching and maintaining good performance, despite its clear relevance (11, 12). This is particularly disappointing given the importance of sensorimotor control and learning in academic attainment (13), neurological deficit (14), and socioemotional development (15).

The possibility of using gameplay to measure skill development has long been recognised (16, 17), allowing the full history of a player’s trajectory to be tracked in an automatic and naturalistic manner (18). Previous research tends to measure skill development in games using holistic measures, such as a player’s score per game (18), proprietary in-game skill metrics (19–21), or multi-player metrics like the ratio of kills and assists to deaths (22). While these provide interesting examples of how skills broadly improve, by representing skill holistically, they do not inform what specific sub-components of behaviour contribute to motor development. For example, an equal improvement in score may arise because of changes in decision-making or motor execution.

Two recent studies have used a commercial FPS aim-trainer to characterise motor performance (23, 24). In FPS games, players see the virtual world from the first-person perspective of their character and use their mouse to control the character’s view, typically to aim towards and shoot enemies. The tasks used are simplified environments that isolate specific patterns of behaviour, such as aiming while shooting stationary targets. The studies have found that broad measures of motor performance, such as hit accuracy and hit rate, improve with practice (24) and that certain features of a movement, such as reaction time and precision of movements, correlate well with a derived measure of motor skill (23).

Similar aim-trainer style games have been embedded into commercial FPS games, finding highly skilled gamers were better on a range of temporal measures (25, 26), and actual gameplay data have been analysed, finding professional players have improved hit rates, lower reaction times, and more efficient movements, compared to amateurs (27).

While the study of skill in FPS games has only recently come to the fore, the act of moving towards a target has been studied extensively in a laboratory setting using visually-guided reaching tasks. Much of our modern understanding of motor control and learning comes from studies where participants interact with a digitising tablet or a robotic arm, representing their movements as those of a cursor moving across a screen, with many recent studies being performed using computer mice or trackpads (28–31). Further, the kinematic analysis of computer mouse movements has been common in the field of human-computer interaction (32–35).

The contingency between onscreen visual feedback and movement is very different between these laboratory reaching tasks and FPS games. Movements of an arm or computer mouse in a typical experiment translate a cursor across a static environment (known in human-computer interaction as Pointing, henceforth *Point*), in much the same way someone would usually interact with their computer’s desktop. In contrast, FPS games use the movement of the mouse to pan and tilt the view of the game while the cursor remains fixed to the centre of the screen, centring peripheral targets in the player’s view (known in the gaming industry as Mouselook, henceforth *Look*). This different feedback resulting from the same movement could illuminate our understanding of motor control because a comparison of sensory feedback is theorized as the primary input determining feedback gains for the control policy during the online control of movements (36). It is therefore possible the feedback differences between contexts could cause people to move in observably different ways. One study did find that Looking took longer than Pointing (37), but movement requirements were not properly equated across contexts. No other work has compared movements between these contexts.

To investigate whether the visual differences between Looking and Pointing lead to observable differences in motor behaviour, we designed a simple task where players had to execute movements in both contexts over the course of the experiment. Participants attempted to make centre-out reach movements to land on and click a target to ‘pop’ it while under imposed time pressure. Spatial and temporal properties of the movements executed in both modes showed high correlation between contexts, with only slightly elevated reaction and correction times in Looking movements. To further explore behaviour in FPS games, we ran a second experiment, modelled on popular aim trainer games, that had participants complete 20 rounds of Looking movements consisting of shots to 48 targets, arranged in seemingly random configurations. Here we observed a range of classic observations from the reaching literature and related several sub-measures of FPS aiming skill to overall performance.

## Results

### Experiment 1

In Experiment 1, participants (n = 50) used the mouse or trackpad of their personal computer to perform a centre-out reach task, where, upon clicking a start-point on every trial, they had to move to and click a target within a time limit to shoot it, otherwise the target would disappear (Figure 1a). The time limit was manipulated throughout a block following a staircase procedure, increasing or decreasing by 30ms in response to an unsuccessful or successful trial respectively, giving participants personalised time pressure that resulted in success on roughly 50% of the trials. Participants completed a block of 320 trials in both the Point and Look contexts, where mouse movements either translated the cursor across a static background or panned and tilted the game’s view while the cursor remained static respectively (Figure 1b). The visuomotor contexts were equated (see methods) so that each required nearly identical mouse movements to shoot the target, such that direct comparisons could be made.

**Figure 1.**
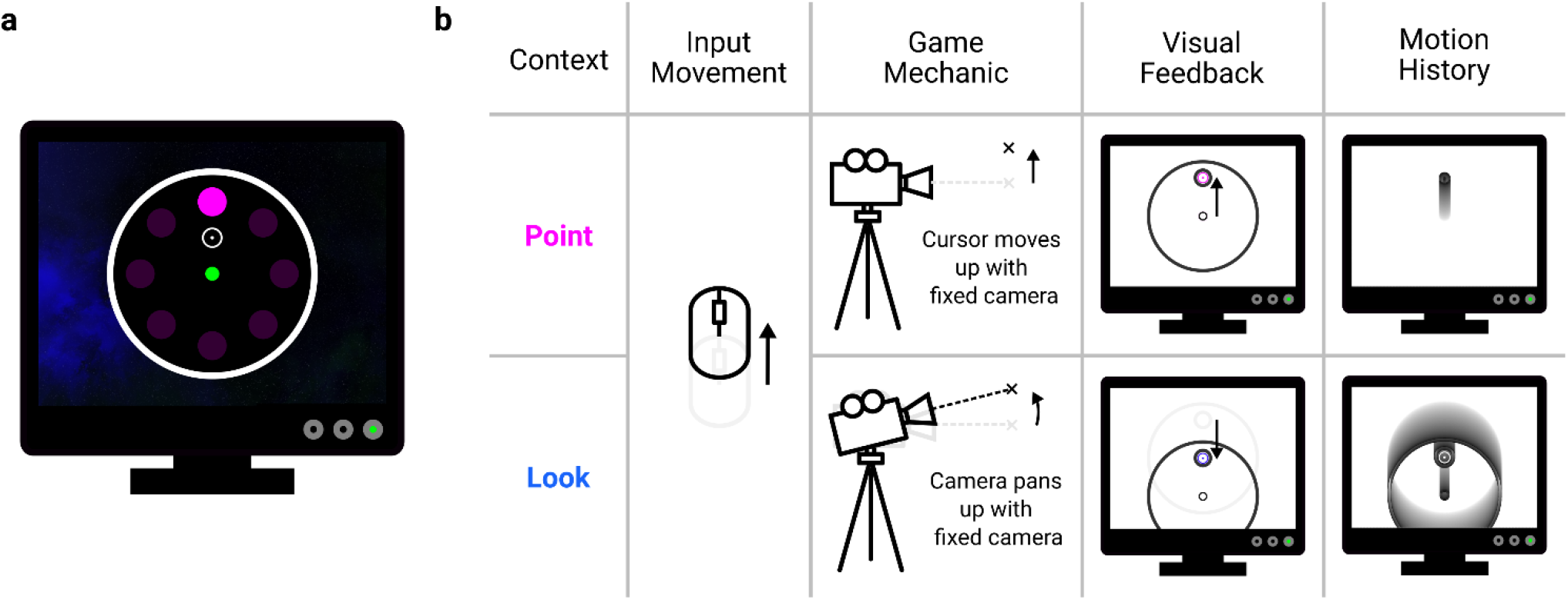
Centre-out reaching paradigm. **(a)** Participants clicked on a start-point located centrally in a circular target plane. Upon a target appearing, located at one of eight potential locations, participants attempted to use their mouse or trackpad to move an FPS-style cursor to the target and left-click to shoot it within a time limit, otherwise it disappeared. **(b)** Participants could control the cursor in two contexts. In the Point context, participants’ mouse movements translated the cursor across an otherwise static scene, typical of most interactions with a computer. In the Look context, participants’ mouse movements panned a camera while the cursor remained central to the camera’s view, typical of FPS style games, giving the visual effect that everything except the cursor translates across the screen. While a common input movement can reach the target in either context, the motion history shows how this difference in feedback leads to different progressions of visual feedback over time.

### Participants required more time to shoot targets in the Look context

Staircase timeouts progressed similarly between contexts, appearing to become roughly asymptotic by the end of the block (Figure 2a). The median timeout over the last 40 trials of each of a high-starting and low-starting staircase showed no significant difference for the Point (t(49) = -0.78, p = .437) or Look context (t(49) = -1.14, p = .259), so we calculated a single asymptotic timeout per context over the last 40 trials of both staircases combined, and used this as our measure of participant skill in this experiment. Asymptotic timeouts per participant were highly correlated between contexts (r(48) = .95, p < .001; Figure 2b), and were longer in the Look context (mean difference [95% confidence intervals] = 53ms [33ms – 73ms], t(49) = 5.18, p < .001; Figure 2c).

**Figure 2.**
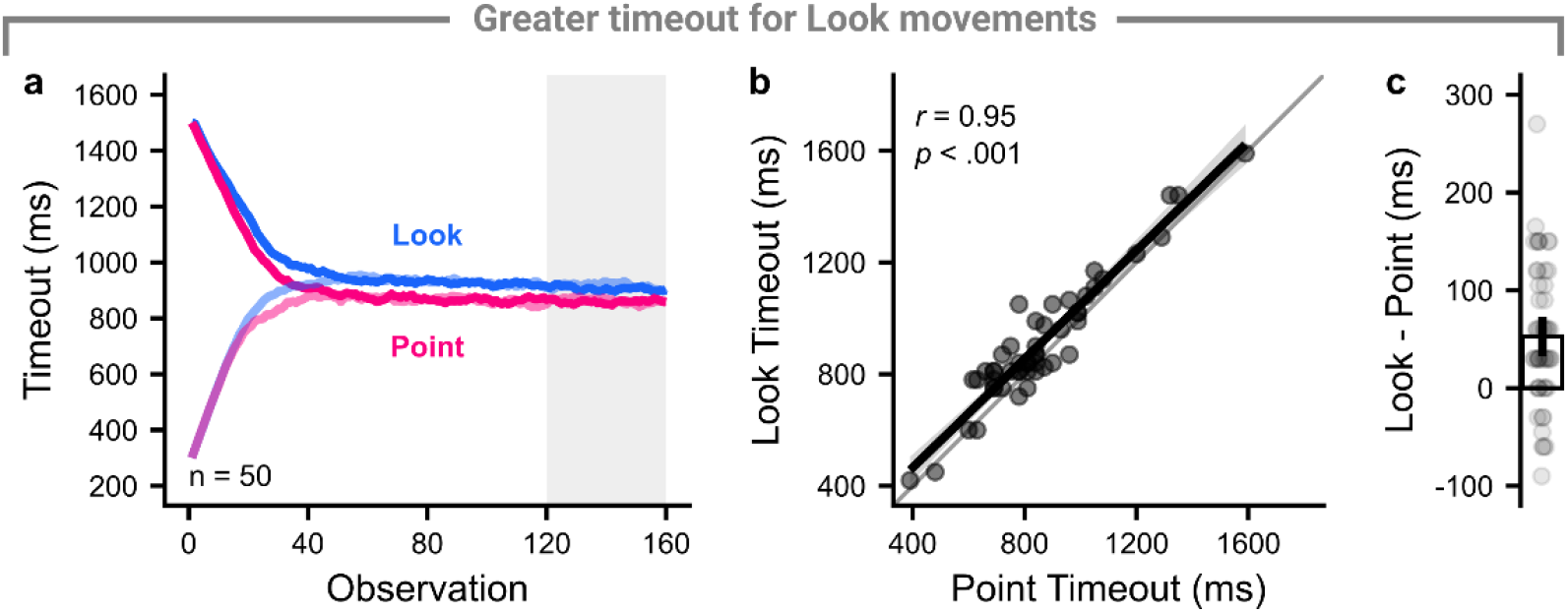
Comparison of trial timeout between contexts. **(a)** Solid lines show the mean timeout across participants per context and staircase, starting either high (darker) or low (lighter). The last 40 trials are highlighted as comparisons between metrics were made over the last 40 trials of each context’s staircases through the rest of this experiment. **(b)** Points show the within-subject median timeout per context, the thick line shows the regression line, and the shaded region shows the 95% confidence interval of the regression line. **(c)** Points show participant differences between the contexts, and the bar and vertical line show the group mean difference and 95% confidence interval respectively.

### Spatial measures are identical between contexts

We used the data within movements to better understand how this time difference arose. On successful trials, participant movements showed variable curvature around the ideal movement path, but average hand paths were reasonably straight and directed to the targets (Figure 3a). Across all participants, average hand paths per target showed little curvature (Figure 3b). While all successful trials end in the target, there were a range of ways a participant could fail a trial. The most common failure (48%) was that participants did not click in time to shoot the target, despite being inside it (Figure 3c-d), but participants also undershot or overshot the target or missed it due to poor directional aim. There appears to be little difference in the hand paths and end-point distribution of movements between Point and Look context.

**Figure 3.**
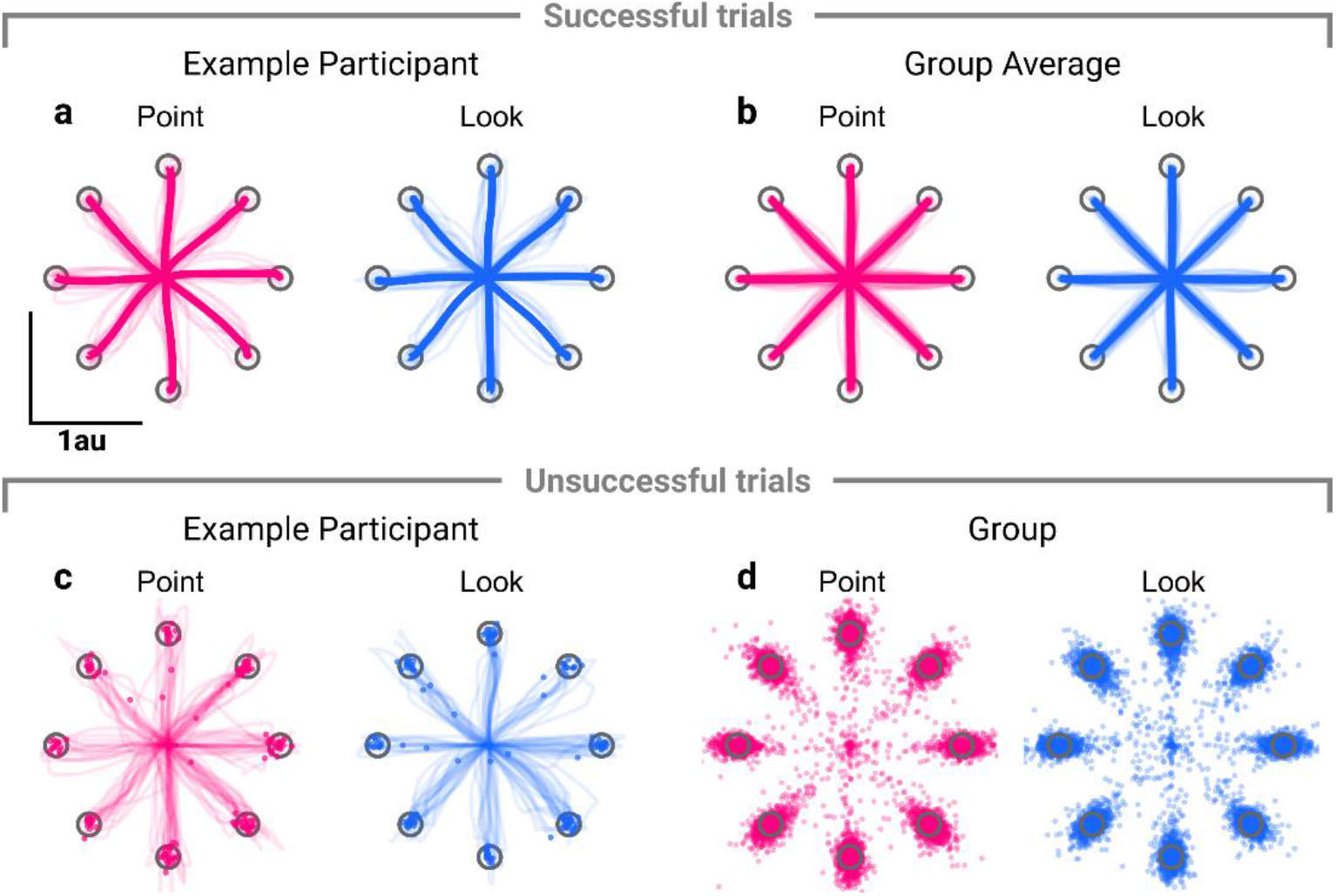
Hand paths for successful trials **(a-b)** and end-point distribution for unsuccessful trials **(c-d). (a)** Hand paths for an example participant on successful trials, both for individual movements (thin lines) and for average movements to each target (thick lines). **(b)** Average hand paths for each target on successful trials, both for participants (thin lines) and for group averages (thick lines). **(c)** Hand paths and end-points for an example participant on unsuccessful trials. Each thin line and point shows an individual trial. **(d)** End-point distribution for all participants on unsuccessful trials.

Finding no apparent difference in hand paths, we investigated four spatial measures that may be able to explain the difference in timeout between contexts. These were the mean and variability of the distance covered at the end of the primary sub-movement (the point at which *discrete* feedback corrections are thought to begin; 38), the variability in angle at the end of the primary sub-movement, and peak speed. Because it was possible for some movement features to be missing on unsuccessful trials, we only analysed successful movements from the last 40 trials of either staircase to ensure a consistent sample across measures.

We assessed both the radial extent over the whole movement and at the end of the primary movement (Figure 4a). The average radial extent over the whole profile showed little difference between contexts (Figure 4b), and the average radial extent at the end of the primary movement was strongly correlated between contexts (r(48) = .81, p < .001), with no significant difference between the Look and Point contexts (−0.01au [-0.03au – 0.01au], t(49) = -1.27, p = .211; Figure 4c). Further, the variability in the distance of the primary movement also correlated well between contexts (r(48) = .63, p < .001), with no significant difference between contexts (−0.01au [-0.02au – 0.01au], t(49) = -1.15, p = .255; Figure 4d).

**Figure 4.**
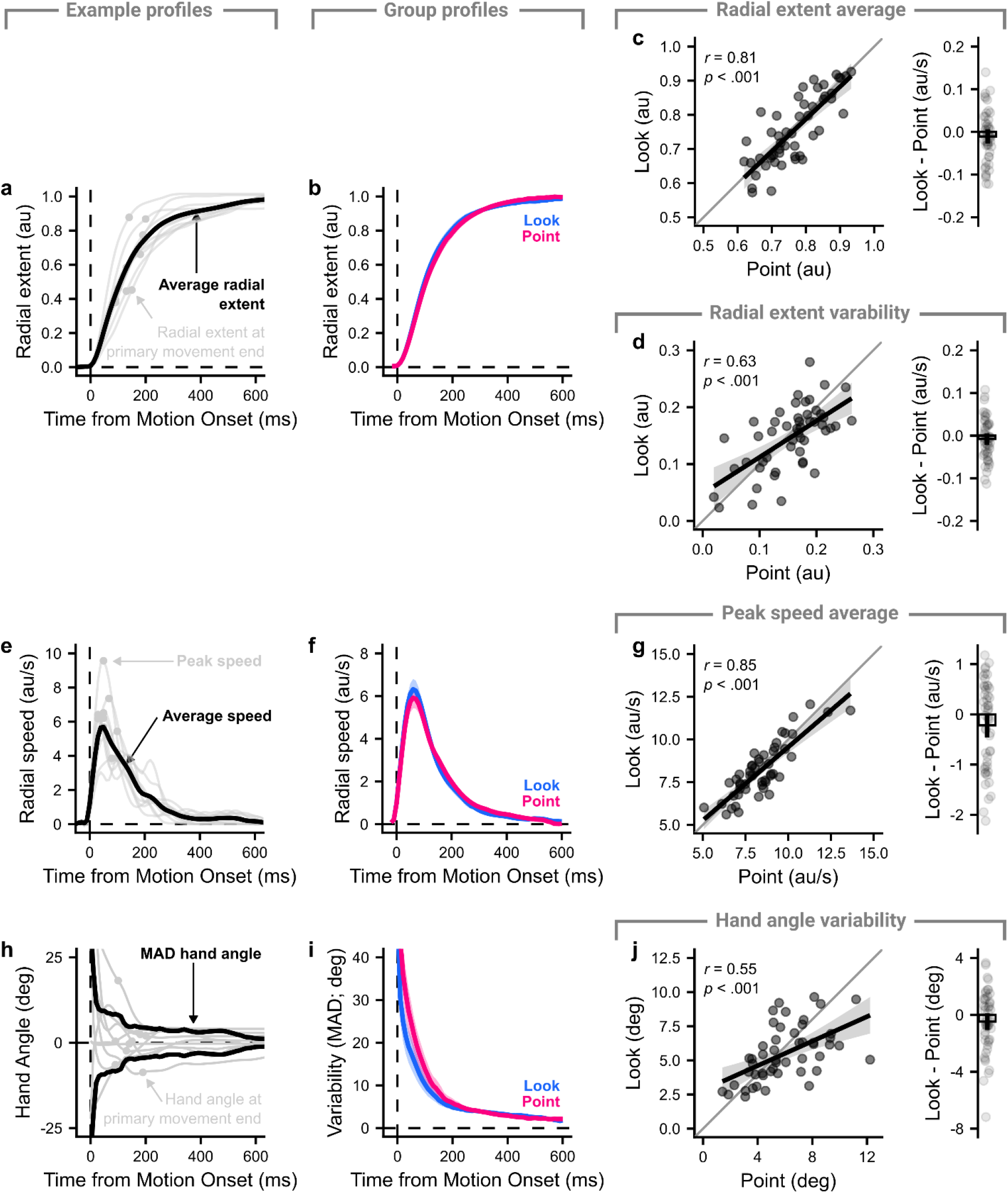
Comparison between contexts on successful trials for derived spatial measures. **(a)** An example of 10 trials showing how the radial extent measures were extracted. The grey lines show the radial extent across time of individual trials, with a point showing the radial extent when the primary movement ended, and the black line shows the average radial extent over the individual trials. **(b)** The lines show the average across participants per context, with the shaded regions showing 95% confidence intervals. **(c)** The average radial extent at the end of the primary movement was compared between contexts. Left panel: Points show individual participants average radial extent, the thick line shows the regression fit and the shaded region shows the 95% confidence interval. Right panel: Within-subject difference in average extent between contexts. Points show participant differences, the bar shows the mean difference and the vertical line shows the 95% confidence interval. **(d)** as **(c)** but with variability in radial extent. **(e-g & h-j)** as **(a-c)** but showing how the radial speed and hand angle were compared between contexts respectively.

A similar analysis was performed for reach speeds (Figure 4e). The average speed profiles for each context were almost identical (Figure 4f), with high correlation (r(48) = .85, p < .001), and no significant difference in average peak speed between the contexts (−0.22au/s [-0.46au/s – 0.02au/s], t(49) = -1.79, p = .080; Figure 4g). Finally, the variability in angle from the target (Figure 4h) was again very similar across the contexts (Figure 4i), with high correlation (r(48) = .55, p < .001) and no significant difference in variability in hand angle at the end of the primary movement between the contexts (−0.48° [-1.07° – 0.11°], t(49) = -1.58, p = .120; Figure 4j).

These results indicate group behaviour is similar across all spatial measures and are therefore unlikely to account for the difference in timeout observed between contexts. The measures may, however, be able to account for good performance overall. All spatial measures were correlated with asymptotic timeout when collapsed across context: average (r(48) = -.63, p < .001) and variability in distance of the primary movement (r(48) = -.50, p < .001), peak speed (r(48) = -.30, p = .032) and variability in hand angle at the end of the primary movement (r(48) = .41, p = .004).

### Temporal measures reveal difference between contexts

Because the spatial measures were all measured after motion onset and prior to corrections, other phases of the movement should account for the context difference. For successful trials, we can measure the acquire time – the total time required to go from seeing a target to shooting it. Acquire time was highly correlated between contexts (r(48) = .96, p < .001; Figure 5a), with a higher average acquire time in the Look context (43ms [26ms – 59ms], t(49) = 5.13, p < .001). Given that the acquire time participants can achieve while being successful determines the asymptotic timeout it is no surprise that the acquire time and asymptotic timeout were nearly perfectly correlated (r(48) = .99, p < .001).

**Figure 5.**
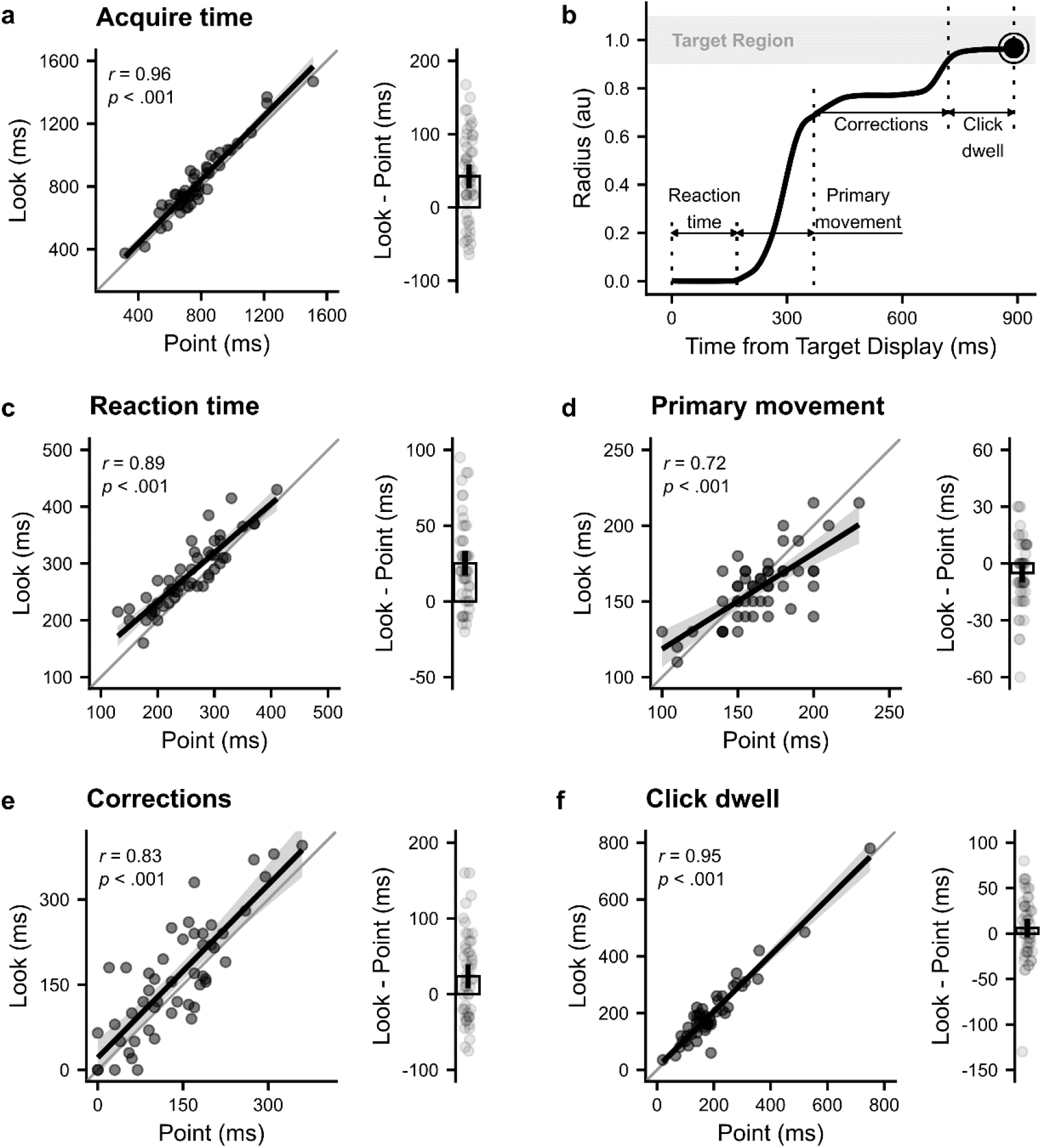
Temporal measures. **(a)** Left panel shows the correlation between acquire time in the Look and Point contexts, with points showing participant medians, thick line showing the regression line, and the shaded region showing 95% confidence interval for the regression line. Right panel shows the difference between the Look and Point contexts, with points showing the within-subject difference, the bar showing the group mean difference, and the vertical line showing the 95% confidence interval in the mean difference. **(b)** Derivation of the temporal metrics. Reaction time: time from stimuli onset to movement onset. Time to peak speed: time from movement onset to peak speed being reached. Online corrections: Time from peak speed being reached to the target being entered. Click dwell: Time from the target being entered to a successful shot. **(c - f)** as **(a)** but for the derived measures.

Using movement kinematics, we can split the acquire time into distinct phases (Figure 5b). We first extracted reaction time, measured as the time from a target being shown to the first sample where the radial speed was above a threshold of 0.5au/s. We also extracted the primary movement time, between movement initiation and the end of the primary movement; the click dwell time, between the target being entered and a successful click being registered; and the correction time, between the primary movement ending and the target being entered. Again, we only use the successful trials from the last 40 observations per staircase.

Across all temporal measures, performance in the Look and Point contexts was highly correlated (Reaction time: r(48) = .89, p < .001, Primary movement: r(48) = .72, p < .001, Corrections: r(48) = .83, p < .001, Click dwell: r(48) = .95, p < .001; Figure 5c - f). Given that these measures combine to give the total acquire time and hence interact with the asymptotic timeout, they should be able to account for the differences between the contexts. Reaction time (25ms [17ms – 33ms], t(49) = 6.24, p < .001) and correction time were significantly greater in the Look context (24ms [8ms – 39ms], t(49) = 2.95, p = .005), but neither the primary movement (−5ms [-10ms – 1ms], t(49) = -1.93, p = .059) or click dwell time (6ms [-4ms – 16ms], t(49) = 1.20, p = .235) significantly differed between contexts (*note that the sum of differences here does not equal the difference in acquire time, as the sum of medians is not equal to the median of sums*). When trials were averaged across both contexts, all measures except for primary movement time were highly correlated with the asymptotic timeout (Reaction time: r(48) = .63, p < .001, Primary Movement: r(48) = -.09, p = .525, Online corrections: r(48) = .76, p < .001, Click dwell: r(48) =.87, p < .001).

### Experiment 2

Experiment 1 showed that kinematic analysis can be successfully applied to FPS-style movements, with task performance decomposed into spatial and temporal metrics that showed high correlations between the Point and Look contexts. Further, all spatial measures correlated with the asymptotic timeout, suggesting they may be important predictors of FPS skill. However, given that this task was designed as a comparison between traditional Pointing tasks and FPS-style mouse Looking, it did not assess kinematic markers of FPS gaming skill within their typical context. We therefore designed a second experiment that only assessed Looking movements in an aim-trainer style task, akin to previous studies (23, 24).

In this experiment, participants (n = 86) completed 20 rounds, each consisting of shots to 48 targets. To start a round, participants clicked a start-point, which made a target appear as a filled circle. The next target location was also simultaneously shown as a hollow, faded circle. For the rest of the round, any time the current target was shot, the next target immediately filled in, and the following target appeared as another faded hollow circle. There was no timeout for movements, but participants were told to complete each round as quickly as they could, with a clock shown during and after the round indicating how long they had taken. The target locations were chosen as a combination of three movement distances (0.4, 0.6, 0.8au) and 8 directions (0 – 315 degrees in 45 degree increments), with each combination sampled twice per round.

### Looking movements show classic reaching observations

To further investigate the similarity between Pointing and Looking movements, we considered whether the Looking movements here showed classic features of reaching movements from the literature, delivered with arm reaches and shown with either actual feedback of one’s hand or with Pointing style feedback.

### Reaches are straight (39, 40)

By arranging movements at their initial position, we can see both at the individual movement level (Figure 6a) and at the group level (Figure 6b), movements are generally straight and aimed directly at the target. The linearity index (maximum perpendicular displacement from a line joining start and end points divided by the length of this line) was calculated for the group, giving an average value of 0.08, similar in magnitude to other observations of horizontal reaching movements (41, 42). This observation is important, as it has been demonstrated that people optimize movements to ensure visual feedback cursors move in a straight line over other factors like metabolic costs (43).

**Figure 6.**
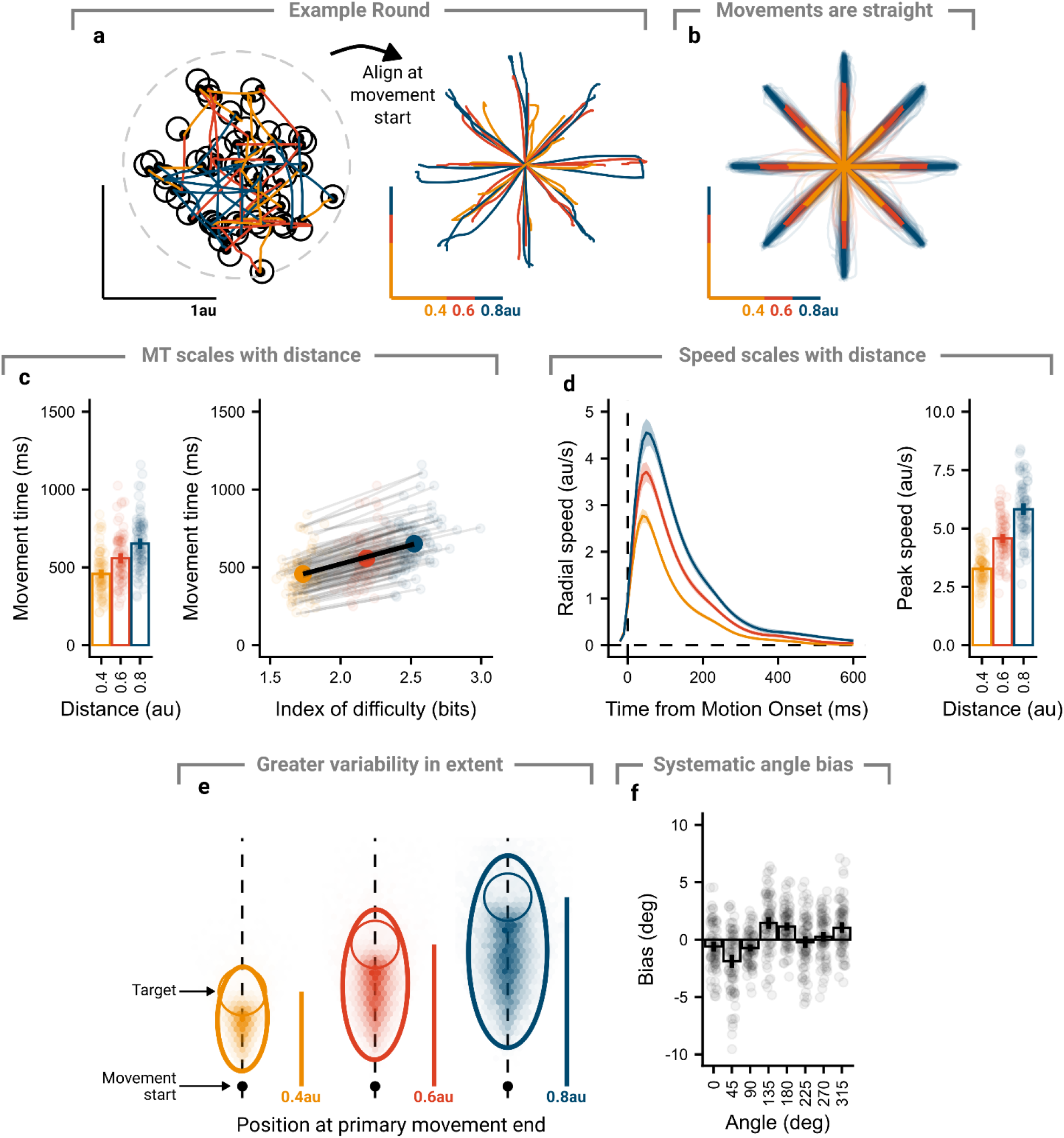
Looking movements show common features of reaching movements. **(a)** Participants completed rounds that consisted of 48 back-to-back movements. While targets had the appearance of being randomly located, they were organised so that all combinations of 3 distances (0.4, 0.6, 0.8 au) and 8 angles (0 – 315° in 45° increments) were tested twice. Arranging movements at their start-points allowed them to be analysed as center-out reach style movements. **(b)** Participant movements were generally straight. Thin lines show participant average movements to each of the 3 distances and 8 directions, with thick lines showing group averages. **(c)** Movement times (from motion onset to successful click) scaled with the target distance. Left panel – points show participant average movement times, bars show group average, and lines show group 95% confidence intervals. Right panel – linear relationship between movement time and index of difficulty, with small points and lines showing participant average and regression slope, and large points and line showing group average and regression slope. **(d)** Speed scaled with movement distance. Left panel – line shows group average speed profile, shaded region shows 95% confidence interval. Right panel - points show participant average movement times, bars show group average, and lines show group 95% confidence intervals. **(e)** Movements showed more variability in extent than direction. Heat map of cursor position at primary movement end, after aligning all movements as if directed to the target directly up. Target shown as a thin circle, 95% error ellipse shown as thick ellipse. **(f)** Primary movements showed a systematic directional bias. Points show participant average directional error, bars show group average, and lines show group 95% confidence intervals.

### Movement time scales with distance (44, 45)

Movement times were found to increase as the required movement distance increased (Figure 6c, left panel). A repeated-measures ANOVA on participant median movement times found a significant main effect of movement distance (F(1.33, 112.84) = 825.37, p < .001), with pairwise comparisons (Bonferroni-Holm corrected) showing greater movement times for greater distances (p’s < .001, d’s > 2.76). Instead of directly looking at the movement distance, the movements could also be compared by the index of difficulty (45), with Fitts’ Law positing a linear relationship between movement time and the index of difficulty. The *effective* index of difficulty was calculated for each participant (46), and group averages taken over participants (Figure 6c, right panel). A mixed-effect model was fit, with a slope for index of difficulty and a random intercept and slope per participant. The population had an intercept of 35ms (p = 0.044) and a slope of 243ms/bit (p < .001), comparable in magnitude to studies of mouse Pointing performance (37, 47).

### Speed scales with distance (41, 44, 48)

Velocity profiles showed scaling with movement distance, where further movements showed higher velocities throughout (Figure 6d, left panel). This can be assessed more directly by finding the peak speed reached on trials, which increased with movement distance (Figure 6d, right panel). A repeated-measures ANOVA on participant median peak speeds found a significant main effect of movement distance (F(1.17, 99.57) = 980.17, p < .001), with greater speeds at greater movement distances (p’s < .001, d’s > 2.66). Further, a repeated-measures ANOVA on the variability of peak speed found a significant main effect of movement distance (F(1.18, 100.52) = 120.10, p < .001), with greater within-subject variability in for greater movement distances (p’s < .001, d’s > 0.86), consistent with signal-dependent noise (49).

### Greater variability along the primary movement axis (48, 50)

To assess variability in feed-forward movement commands, we evaluated early kinematic markers, which have been shown to correlate well with end-point kinematics for uncorrected movements (50). We used the location at the end of the primary movement to assess variability. Cursor positions at the primary movement end were fit with 95% error ellipses per participant (Figure 6e). A repeated-measures ANOVA on participant ellipsoid aspect ratios found a significant main effect of movement distance (F(1.93, 164.03) = 52.26, p < .001). Aspect ratios were significantly greater than 1 for all distances (2.14 – 2.51, p’s < .001, d’s > 2.16), indicating more variability in extent than direction (48, 50), and increased with movement distance (p’s < .001, d’s> 0.41).

### Systematic angle bias (51, 52)

The group showed a consistent bias in the angle of their movement at the end of the primary movement. The average directional error appeared to follow a curve with two peaks and troughs, consistent with previous studies using arm reaches and mouse Pointing (52, 53), as well as studies in progress from our laboratory on mouse Pointing movements, though the pattern of bias appears to vary based on a variety of factors like vision and posture. A repeated-measures ANOVA on participant median directional errors showed a main effect of movement angle (F(4.46, 334.29) = 25.16, p < .001).

### Concurrent improvements in temporal and kinematic variables

In line with a previous study looking at effects of practice in an aim-training game (24), participants improved their performance over the course of the experiment, here operationalised as improvements in acquire time (Figure 7a). Over the twenty rounds, participants improved their acquire time by 317ms ([249 – 385], t(85) = 9.17, p < .001, d = 0.99). The improvements appeared to consist of two phases, with a rapid improvement (75% of total gain) over the first 5 rounds followed by a sustained slower improvement through to the end of the experiment.

**Figure 7.**
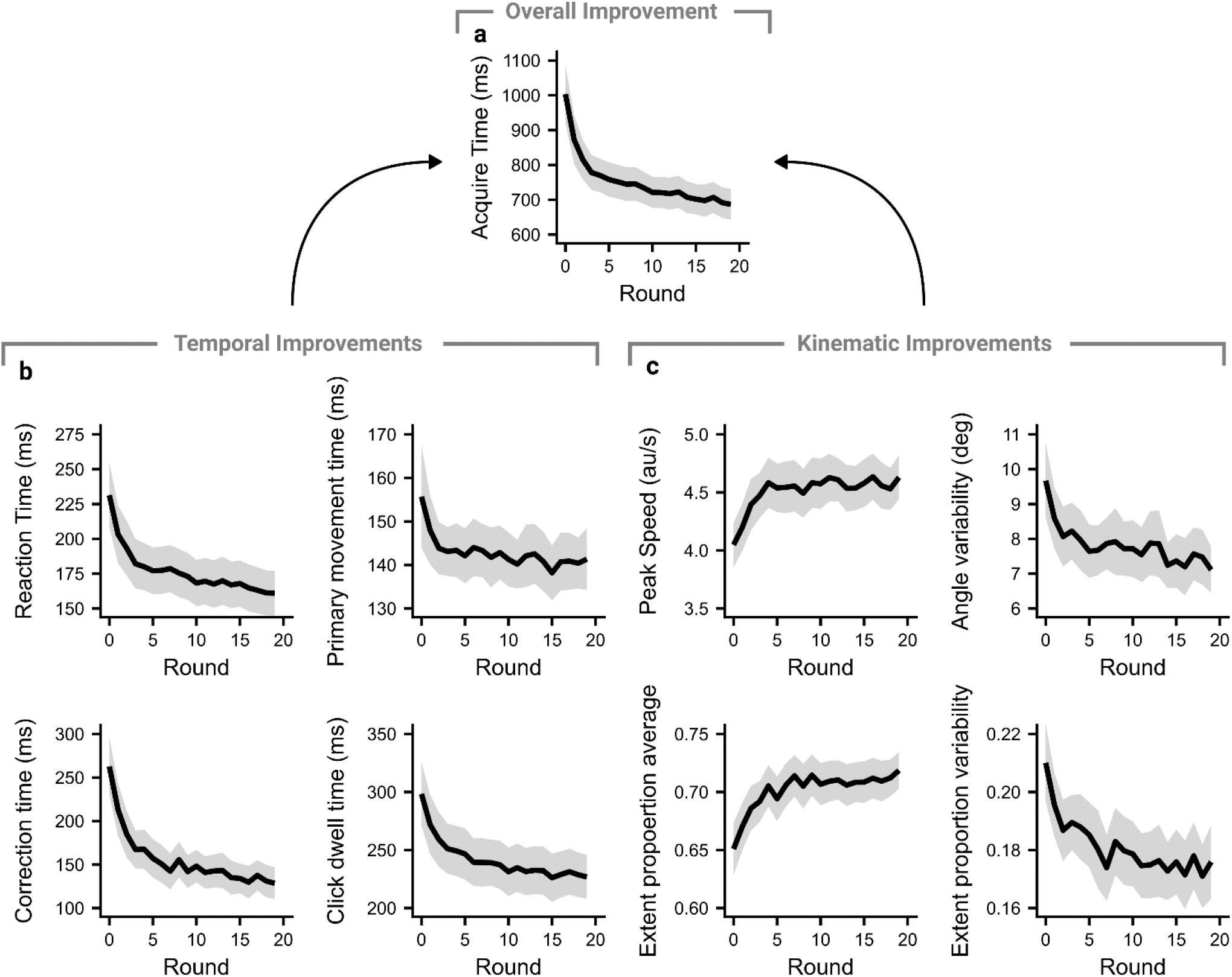
Temporal and kinematic variables show concurrent improvements over the experiment. **(a)** The main measure of performance on the task, the acquire time, improved continuously over the course of the experiment. The improvements appeared to consist of two phases – an initial rapid improvement, followed by a sustained but slower improvement. **(b)** The constituent temporal variables all show improvements over the experiment. **(c)** The tested kinematic variables also show improvements over the experiment. For all panels, thick line shows group mean and shaded regions show 95% confidence intervals.

As in the first experiment, we can divide the total acquire time up into different phases of the movement (Figure 7b). All phases of the movement showed significant improvements (Reaction time: 70ms [55ms – 86ms], t(85) = 9.03, p < .001, d = 0.97; Primary Movement: 14ms [4ms – 25ms], t(85) = 2.67, p = .009, d = 0.29; Corrections: 134ms [105ms – 164ms], t(85) = 8.92, p < .001, d = 0.96; Click dwell: 72ms [51ms – 92ms], t(85) = 6.98, p < .001, d = 0.75). Given that each phase of the movement improved, we would expect this to be accompanied by improvements in key kinematic variables that might determine the duration of certain phases. All four kinematic variables showed significant improvements over the experiment (Peak speed: 0.58au/s [0.41au/s – 0.76au/s], t(85) = 6.51, p < .001, d = 0.70; Angle variability: 2.56° [1.60° – 3.52°], t(85) = 5.25, p < .001, d = 0.57; Extent average: 0.07 [0.05 – 0.09], t(85) = 6.68, p < .001, d = 0.72; Extent variability: 0.03 [0.02 – 0.05], t(85) = 4.37, p < .001, d = 0.47).

### Kinematic variables explain individual differences

As in Experiment 1, we wanted to understand whether these kinematic variables were important predictors of participant performance. To do this, we looked at whether the acquire time by the end of the experiment (the last round) was significantly predicted by any of the kinematic variables shown in Figure 7c. A linear model with acquire time as the dependent variable and the four kinematic measures as independent variables was fit. The model (F(4, 81) = 16.06, p < .001, adjusted R^2^ = 0.41) showed the average and variability of primary movement extent were significant predictors of acquire time across participants, with a 1SD increase in either measure leading to a 70ms decrease (p = .001) and 114ms increase (p < .001) in acquire time respectively. Neither peak speed (p = .207) nor variability in angle from the ideal path (p = .373) significantly predicted acquire time. That such a small number of kinematic variables can explain so much of the variance in performance between participants highlights the utility in trying to understand the subcomponents of skilled FPS movements.

## Discussion

Despite widespread interest in the cognitive and perceptual abilities of video gamers, only recently have studies begun to assess the motor skills demonstrated in these games. We aimed to isolate the skill of aiming at and shooting targets and analyse its sub-components using the kinematics of participant’s movements. We found movements in an FPS-style Look context to be nearly identical to those in a more traditional Point context across a range of spatial and temporal measures, only differing slightly in reaction and correction times. We also showed that FPS movements are consistent with several classic observations from the reaching literature. We observed concurrent improvements in all tested spatial and temporal metrics with practice and found that the participant’s average and variability in the extent of primary movements accounted for just under half of the between-participant variance in task performance. Our metrics can be generalised to studying FPS movements in other tasks, and actual gameplay data, providing an opportunity to advance our understanding of what factors set the best players apart.

### Studying skill in first-person shooter games

Typical approaches to the study of skill involve abstract laboratory tasks or making restricted observations of real-world behaviour using specialised equipment. Video games provide a unique opportunity to observe the development and maintenance of skill in its natural environment by simply recording the game inputs, typically provided by a keyboard and computer mouse. Only recently have studies begun to leverage this to assess skilled motor behaviour in FPS games (23–27), though earlier studies have taken this approach to understand the skill in real-time strategy games (20, 54, 55).

We found that participants improved their Look performance with practice during the aim-training task (Experiment 2), reducing the total time to acquire targets by around 300ms. This improvement appeared to proceed in two stages – an initial rapid learning where the majority of the performance gains were realised, followed by a sustained but slower learning. This dual-rate pattern is consistent with improvements in hit speeds previously found (24), and with classic studies of motor learning (56–58). To move beyond broad measures of performance, we used the kinematics of participants’ movements to understand this development across a range of spatial and temporal metrics, finding that players improved across all metrics tested. Further, we characterised the importance of these measures to overall performance, finding the average and variability in distance covered by the end of the primary movement to be significant predictors.

While other studies have addressed similar questions and observed results consistent with our own, finding that similar temporal measures improved with practice (26) and reaction time and movement precision were significant correlates of motor skill (23), we believe our metrics have a number of advantages over those previously reported. Where Toth et al. (26) investigated an analogous measure to our click dwell time (verification phase), it was based on when participants’ movement speed last dropped below a velocity threshold. This method permits the verification phase to begin before the target is reached, as sub-threshold movements continue to traverse the remaining distance. We instead delineated correction and click dwell phases by when the target was entered for the last time, ensuring that our click dwell metrics only accrued time when a trial could have been ended successfully. Further, Donovan et al. (23) characterised movements as being composed of sequences of distinct sub-movements and pauses, and derived metrics from sigmoids fit to each sub-movement. This forces a model onto the data that assumes that sub-movements are distinct and ballistic, yet individual trial speed profiles shown in Figure 4e and other studies investigating speeded reaches with online feedback (59–62) find asymmetric speed profiles and discrete corrections while speed was still high. Therefore, we used well-validated techniques to identify discontinuities in speed profiles (63, 64) to characterise spatial metrics at the end of the primary movement, making fewer assumptions about the exact form of the movements.

While the studies discussed thus far, and our own, utilised purposely simplified aim-trainer style tasks (to isolate specific components of behaviour), the kinematic analyses described here are also applicable to FPS games generally. Indeed, similar analyses have been performed on actual gameplay data (27), finding professional players execute more effective shots, exhibit lower reaction times, and make more efficient movements. We believe that the utility of this approach will be most apparent if researchers can collect such gameplay data at scale to describe differences within a wide population of players rather than between groups of players of different experience. However, as the environmental constraints are weaker than those in an aim trainer (like non-stationary targets, more than one ideal aiming location, and simultaneous character movement), the analysis techniques will need to account for the greater range of behaviours that may be observed.

### First-person shooter games inform our understanding of the neural control of movement

To understand whether there were any fundamental differences in how participants controlled their movements in the Look and Point contexts, we fully equated the movements required in each context such that an identical mouse input would produce the same relative movement in the game. Participants required a 53ms larger time limit to complete Looking movements, with our kinematic decomposition indicating the difference was accounted for by larger reaction and correction times, each taking around 25ms longer, while spatial features of the movements were nearly identical between contexts.

The only other study to compare these two contexts found Looking movements took around 230ms longer than Pointing movements (37). This difference was quantified between the intercepts of linear regressions of movement time against index of difficulty, sometimes interpreted as measuring processes distinct from the movement itself (65), so could be consistent with our finding of greater reaction times. However, Pointing was assessed via cursor movements to 2D rectangular targets, and Looking in Unreal Tournament where users shot alien avatars, so several factors beyond the visual feedback could contribute to differences found between these tasks, for example if there were greater perceptual demands to locate targets in the 3D task. Such confounds hamper interpretation of the observed differences in Looser et al. (37), but are addressed by our task design, which shows more modest context differences.

Despite the two feedback conditions entailing fundamentally different mappings between movement and visual motion feedback, behaviour was highly similar. This is important because preeminent models of motor control propose that sensorimotor behaviour is guided by a sensory state estimate generated between a continuous comparison of actual sensory feedback and a forward model’s predictions of the sensory consequences of motor commands (66, 67). Each movement context in these computational frameworks has its own internal model, which entails forward and inverse models (68, 69). Critically, if forward models predict primary sensory input, then feedback control would require different forward models for Point and Look conditions, as the sensory consequences of the same motor command evolve very differently over the course of a movement.

It is unlikely that Look and Point contexts require entirely different internal models. Most of our participants (38 of the 50) reported not playing FPS games and 11 reported playing no video games at all. All were regular computer users, with a wealth of experience in the Point context. If distinct internal models were required for these conditions, the well-practiced Point movements should be relatively accurate and fast while Look movements would be slow and inaccurate, requiring *de novo* development of an internal model or association of arbitrary sensory feedback with existing motor commands. Visuomotor tasks that require such learning show initial poor performance that requires an extended period of practice to overcome (58, 70–73). Instead, the majority of participants in our experiments could readily perform movements in the Look context and their performance was highly correlated across the two conditions.

Our results suggest that participants’ movements were largely impervious to the specifics of visual motion that result from movement. Instead, they appear to infer the relative positions of target and effector despite large differences in visual motion. This observation can fit within existing motor control theory if forward models in these frameworks (74, 75) make predictions of displacement vectors for visuospatial coordinates, rather than predictions of primary sensory feedback. This would allow the motor system to use the same internal model for both movement contexts, without a requirement to specify the categorical differences in visual motion (or more qualitative factors such as colour). These complex facets of a visual scene would, however, need to be parsed upstream in higher visual areas.

There is good evidence that the posterior parietal cortex combines hand and gaze reference frames into a hand-target displacement vector that is used to plan and correct movements (76, 77). We propose that visual target and effector positions for Look and Point would be parsed by the visual system at or before this step in the parietal cortex, and that forward model sensory predictions would be at the level of the combined hand-target displacement vector rather than any primary sensory modality. The ability of such a modular hierarchy to deal with stark differences in visual motion may help explain why movement vectors have been shown to be a critical link for both planning and motor learning (48, 78– 80), and why visuomotor adaptation may occur at the step between parietal cortex and premotor cortices (81).

In pointing movements, the eyes will typically saccade to the movement target before the hand movement is initiated and fixate it for the duration of the movement. Larger hand movement errors are made if the target is not foveated during these movements, suggesting this process improves target localisation (82, 83). As both contexts appear identical before movement is initiated, we might expect a similar initial saccade for both Point and Look. However, upon moving the mouse, the target will begin to move in the Look context, which might require an extra saccade or smooth pursuit to maintain foveation during the final parts of the movement, with previous work suggesting the majority of fixations in FPS games are located around the aiming reticle (84). Future work, where the patterns of eye movements are compared between the two contexts, should provide clarity on whether this is indeed the case, and if so whether it can account for increased correction times in the Look context.

### Wider applications

A perennial issue for FPS games is detecting cheats who gain an unfair advantage through use of third-party software that allows, for example, enemies to be seen through walls or automatically aimed at. Analysing gameplay data for abnormal aiming abilities or other behavioural patterns has been suggested as an effective method for cheat detection (85–87). Our analysis of kinematics is both more general and precise than previous work, building on decades of kinematic analysis of pointing movements. We believe our analysis could be readily adapted to help detect cheaters.

As professional esports teams increasingly look to use analytical approaches to inform training programmes (88), the decomposition of a player’s movements into kinematic metrics may allow tailored training on certain aspects of a movement. For instance, a coach could increase focus on clicking predictively to reduce click dwell time. Further, players in teams are typically assigned a specific role to fulfil, like executing accurate shots with a sniper rifle, and a more thorough understanding of the strengths and weaknesses of the team roster may allow better role assignment.

## Methods

### Participants

Participants were recruited through the online testing platform Prolific and were paid £6 upon completion. They were only recruited if they resided in the UK or USA, had English as a first language, and had a Prolific approval rating of 95% or above. Given the experiments were completed online, which is typically associated with noisier responses within and between participants (e.g. Tsay et al., 2021), we applied stringent screening criteria to ensure the final sample was sufficiently high quality. The experiments were approved by the School of Psychology Ethics Committee at the University of Leeds, and participants gave informed consent via a web form prior to starting the study.

### Experiment 1

12 of the 62 participants were removed from analysis. We removed 5 participants whose frame rate was either too low (<30fps, as kinematic analysis resolution was poor for low frame rates) or differed between contexts (>10fps, to remove frame-rate dependent changes in performance). A further 3 participants were excluded because their performance had not stabilised by the end of either block (success rate more than 10% away from intended 50% success rate over the last 40 trials of both staircases per context). Finally, 4 participants were excluded whose difference in timeout between the two contexts was above 3 median absolute deviations (MADs; calculated throughout using a consistency constant of 1.4826 to make it a robust estimator of standard deviation) away from the group median difference (between 4.72 – 9.78), representing a subset of participants who struggled to perform in the Look context. Their performance was around 500-1000ms worse in the Look context, with greatly increased reaction and correction times. The latter appeared to be driven by shorter primary movements (∼0.3au reduction), requiring greater feedback corrections, which introduced weak significant differences between contexts for the primary movement extent and peak speed. While the results were otherwise the same at the group level with their inclusion, with strong correlations between contexts, and would not change our interpretation of the results, we chose to exclude these participants so that the group-level estimates of context differences were not unduly influenced by a small number of outliers. This gave a final sample of 50 participants (16 male, 34 female; mean age ± SD = 38 ± 13, age range = 21 – 66).

### Experiment 2

14 of the 100 participants were removed from analysis. We removed 10 participants whose average frame rate was less than 30fps or dropped by more than 10fps between the first and last round. A further 4 participants were excluded whose median acquire time was more than 3 MADs from the group median over the experiment (3.07 – 4.92). Exclusion of these participants does not change any statistical results. This gave a final sample of 86 participants (44 male, 41 female; mean age ± SD = 43 ± 13, age range 22 – 78).

### Apparatus

Participants used their own personal computer to complete the experiments and were restricted to users on a laptop or desktop using Prolific’s screening tool. The experiment was created using the Unity game engine (2019.4.15f) and the Unity Experiment Framework (89), and delivered via a WebGL build hosted on a web page, with data uploaded to a remote database. Given participants used their own computers with varying sizes and aspect ratios, the physical size of the task was not consistent across participants but was developed to be visible on a 4:3 aspect ratio monitor. The height of the scene, 4 arbitrary Unity units (au) always took up the full height of the participant’s monitor, with wider aspect ratios featuring more of a task irrelevant background texture. Full screen was forced throughout, and during the experiment the desktop cursor was hidden and locked so it could not be used to interact with the web page. Instead, the raw mouse or trackpad input was used to perform in-game movements of the cursor (eliminating mouse acceleration from the operating system). The sensitivity of in-game movements was initially calibrated to be similar to the participants’ desktop cursor, which could be adjusted in a calibration stage.

### Equating pointing and looking movements

Participants used their mouse or trackpad to manipulate the in-game cursor position in either the Point or Look contexts. A key requirement for the experiments were that a particular target could be reached in either context using an identical mouse input. For the Point context, the experiment is set up so that a participant’s view of the scene is provided by a static orthographic camera. The raw mouse input on a given frame, provided as a difference in mouse position from the previous frame, is multiplied by the participant’s calibrated sensitivity and added to the current cursor position. This setup mimics a typical 2D reaching task.

For the Look context, the raw mouse input is instead used to rotate an orthographic camera about a pivot, while the cursor is fixed central to the camera’s view. For a given translation of the cursor from the middle of the workspace along a single axis in the Point context, *o*, this can be formulated as the opposite length of a right-angle triangle, which using a basic trigonometric identity gives *o* = *a* × tan(*θ*), where *a* is the distance between the rotation pivot and the plane on which the cursor should translate and *θ* is the required rotation. For small rotations, a small-angle approximation holds where *tan*(*θ*) ≈ *θ* and therefore *θ* ∝ *o*. The task was developed to have a maximum workspace of 1au in any direction from the centre, and a distance of 50au between the camera’s rotation pivot and the target plane was used to ensure only small rotations were required. We then simply need to multiply the input by a coefficient, *c, to get* a rotation that approximates the equivalent Point translation. Here we pick a coefficient that minimises error at the target distance, 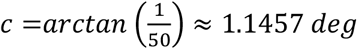. *Over t*he intended workspace, this gave a maximum difference between the cursor’s position on the target plane in the Point and Look modes of 5.13×10^−5^au, which equates to a sub-pixel difference on a 1080p monitor. The raw mouse input on every frame was therefore multiplied both by the participant’s calibrated sensitivity and this rotation coefficient, and added to the camera’s current rotation. This provides a general framework for building experiments where traditional mouse Pointing or FPS-style mouse Looking can be used and compared in the same task.

### Experimental task and procedure

In both experiments, participants first filled in details on their age, gender, whether they were using a mouse or trackpad, a desktop or laptop, and then pressed a button to ensure they could hear game audio. During this stage participants used their usual desktop cursor to navigate the form, and the cursor movements in pixels and Unity game units were tracked to provide an initial calibration for the in-game cursor sensitivity that was around their regular desktop cursor sensitivity. Following this, the desktop cursor was hidden and locked, and interactions were only possible using their in-game cursor.

Participants were shown a brief cut-scene to provide exposition for the game (popping deadly non-sentient space-bubbles), before progressing to a tutorial that introduced elements of a trial sequentially and interactively. At the end of the tutorial participants were required to complete practice trials in each context, during which they could toggle context and adjust their cursor sensitivity. After completing at least 20 trials in each context, they could progress to the main task.

### Experiment 1

Participants used their computer mouse or trackpad to try to move to and shoot a target using an FPS-style cursor in either the Look or Point mode, depending on experimental condition. Participants saw a circular target plane (3au diameter), on top of which targets could appear. The target plane was black, with a thin white ring around it. Behind this was a dark, space-themed background. A start-point (0.1au diameter circle, initially coloured orange) was located at the centre of the target plane.

Participants were required to move their in-game cursor, a small white circle surrounded by a thin white ring (diameter 0.15au), to the start-point and left-click to initiate a trial. If participants were within 0.05au of the start-point when making a homing movement, the cursor snapped to the start-point to provide an FPS-style “auto-aim”. Upon left-clicking the start-point, the colour of the start-point would turn green to indicate participants could move and a target (0.2au diameter magenta circle) would immediately appear on the target plane in one of eight locations. Potential targets were placed in increments of 45° around a virtual circle of radius 1au, where 0° represented a target directly right of centre. Participants had to move to and click on the target within a time-limit to shoot it, in which case it exploded and a shooting sound was made, otherwise it would disappear and make a whooshing sound. A shot was only successful if, at the time of a left click, the centre of the cursor intercepted the target circle. This feedback was provided for 300ms, during which the start-point was coloured grey. After the feedback disappeared, the start-point turned orange, indicating a new trial could begin.

Participants completed 640 experimental trials, split into a 320 trial block of Pointing and a 320 trial block of Looking. The order of the blocks was counterbalanced across participants, with half completing the Point block first. The blocks were arranged into cycles of 8 trials, where each of the possible target angles was tested once in a random order. To induce the sort of time pressure experienced in a typical FPS game, the time limit within which targets had to be shot was continuously staircased throughout the experiment, using a 1-up / 1-down method. For each block two staircases were interleaved, one ascending (starting at a time limit of 300ms) and another descending (starting at 1500ms). After a successful or unsuccessful trial, the time limit was reduced or increased by 30ms respectively. By the end of each block, the time limit should become asymptotic at a value that gives roughly a 50% success rate. Participants were given a self-paced break every 80 trials, and between blocks where participants were informed the context would switch. Following completion of the experimental trials, participants completed a questionnaire probing their enjoyment of the game, perceived lag in the game, gaming experience, and any other comments.

### Experiment 2

Participants only completed movements in the Look context. Experimental trials began the same as in Experiment 1, up until participants had clicked the start-point. After the start-point was clicked, it disappeared and a target appeared at one of three distances (0.4au, 0.6au, 0.8au) and eight angles (0° to 315° in 45° increments). Simultaneously, the upcoming target after the current had been shot was shown as a hollow, faded magenta circle. This was implemented as pilot testing showed without knowledge of the upcoming target, participants moved towards the workspace centre following a successful shot, presumably to be equidistant from any given workspace area until the new target had been visually processed. Upon successfully shooting the current target, the upcoming target immediately became the new current target, and a new upcoming target was revealed. The new upcoming target was always located in a new location at one of the three distances and eight angles away from the current target. Participants continued shooting targets until 48 targets had been shot. While movements had no time limit, participants were told to complete each round as quickly as possible, with an on-screen timer visible during and after the round showing how long it had taken.

The 48 targets per round were arranged into two uninterrupted cycles, where each combination of the three distances and eight angles were tested once per cycle, with the criteria that no individual target could be located more than 1au from the centre of the workspace. The position of targets in each round was generated by simulating new sequences until this criteria was met. This had the effect of making seemingly random sequences of targets, while controlling the statistics of the reaches and staying within the intended workspace. Participants completed 20 rounds of 48 movements and were given a self-paced break between each round. Following completion of the rounds, participants completed a questionnaire probing game enjoyment, perceived lag, strategies they used to improve, gaming experiment, and any other comments, with many participants indicating they’d enjoyed playing this experiment.

### Data analysis

Data were processed using R (version 4.2.2). The cursor position time series data, sampled at the participant’s refresh rate by Unity, was resampled to a standard frequency of 100Hz using linear interpolation. This resampled data was then filtered using a second order, low-pass Butterworth filter with a 15Hz cut-off in the forward and reverse directions to give zero lag, with the start and end of each trial’s time series temporarily padded to remove transient effects of the filter. Movements were segmented in Experiment 1 by extracting the period between the start-point being clicked and an outcome being registered, and in Experiment 2 by extracting the period between the previous and current target being clicked. Velocity, acceleration, and jerk in the radial direction were obtained by numerically differentiating the position data in polar coordinates (using *pracma’s* gradient function), centred on the cursor position at the start of each movement segment. To visualise hand paths, movements in cartesian coordinates were resampled into 100 evenly spaced points between the start and end of the movement segment. Participant averages were produced as the mean position at each interpolated time step, and group averages found by averaging over all participants. The hand paths were used to derive spatial and temporal metrics, with within-subject averages calculated using medians to account for the typically skewed distributions of temporal measures.

### Staircasing metric

In Experiment 1, the continuous staircasing led to participants being successful on roughly half the trials by the end of a block. We extracted each participant’s median timeout over the last 40 trials of each staircase per block to quantify the time required to be 50% likely to execute a successful shot, as this region appeared to contain roughly asymptotic behaviour.

### Spatial metrics

Four spatial metrics were focussed on through both experiments, similar to metrics previously used (23). The progression of each measure could be visualised across a trial, by aligning movements at motion onset and calculating the relevant measure per time-step. However, this method produces profiles that do not necessarily reflect the individual trials well (e.g. in Figure 4e, the average speed profile looks smooth whereas many individual trials show secondary speed peaks), so measures were also extracted at a single, theoretically meaningful point in time. In Experiment 1, participant summary measures were calculated over successful trials from the last 40 observations per staircase in each context to ensure a consistent sample, as unsuccessful trials may be missing some movement features. In Experiment 2, measures were calculated over all movements within each round, giving one observation of each measure per round per participant.

#### Average Peak speed

The peak speed was the maximum radial speed reached on the trial, and medians were taken over these values to find the average peak speed reached.

#### Average extent at the end of the primary movement

For each time step, the extent in the direction of the current target was found by comparing the current cursor position to the straight line joining the cursor position at the start of the movement and the centre of the current target (ideal path). The current distance from the movement start point, and the angle from the ideal path, could then be used to find the extent in the direction of the target using *r × cos (θ)*. This could be converted to a proportion by dividing by the length of the straight line between cursor start point and target centre, which was useful for Experiment 2 to collapse across movement distances.

We were particularly interested in the average extent at the end of the primary movement, as this is purported to represent where discrete feedback corrections begin (38), and hence should determine the magnitude of the feedback correction required. To identify the end of a primary movement, we followed a similar procedure to other papers (63, 64). Following the peak radial speed, we looked for any timestamps where (a) the radial speed fell below the movement threshold, indicating the movement either terminated or was about to reverse direction, (b) the radial acceleration crossed from negative to positive, indicating participants had sped up, or (c) the radial jerk crossed from positive to negative, indicating participants were ‘braking’. The proportion of the extent covered was then found at the timestamp where the primary movement ended, and the median was taken to find the average in this value.

#### Variability in extent at the end of the primary movement

The variability was found by calculating the MAD of the proportion of the extent covered at the end of the primary movement.

#### Variability in angle at the end of the primary movement

We also extracted the hand angle, the difference in angle between straight lines connecting the starting point of the movement to either the target (ideal path) and the current cursor position. Positive hand angles indicated counter-clockwise errors. The MAD was calculated over hand angles at the end of the primary movement.

### Temporal metrics

The kinematics were also used to establish temporal variables that sum to the total time required to shoot a given target. Reaction time was calculated first, here defined as the difference in time between a current target being shown and the first time where radial speed rose above 0.5au/s. The remaining period of the trial could be split up into further phases. The time between motion starting and the primary movement ending gives the primary movement time, and the time between the cursor entering the target and a successful click being registered gives the click dwell time. If the end of a primary movement occurred after the target had been entered, it was shortened to end when the target had been entered, to make the primary movement and click dwell phases mutually exclusive. The remaining portion, between the primary movement ending and the target being entered, can then be treated as the correction time, the time required to execute discrete corrections to bring the cursor within the target. These phases are marked on a single example trial in Figure 5b. Average metrics were produced per participant by taking medians over the successful trials from the last 40 observations of each staircase per context in Experiment 1, and over all movements per round in Experiment 2.

### Classic metrics

In Experiment 2, we also compared the movements to classic observations from reaching tasks, where additional metrics needed calculating. To understand movement time scaling with distance, the median movement time was calculated per distance, and also compared across effective index of difficulty, calculated following standardised methods (46). Effective index of difficulty was calculated as *ID = log*_*2*_*(1 + D*_*e*_*/W*_*e*_), where *D*_*e*_ is the average straight-line distance between movement start and end points, and *W*_*e*_ = 4.133*σ*, where σ was the standard deviation of the end-points along the ideal direction of movement. This was calculated per nominal target distance per participant. Peak speed scaling was assessed by averaging the speed profiles or peak radial speed per target distance, or calculating the MAD of peak speed. Aiming biases were found by calculating the median hand angle from the ideal path per target angle, after hand angles outside 3 MADs from the group median were removed. The comparison of variability in extent and direction was performed by aligning movements at their start-point, rotating movements as if directed to a 90° target, and isolating the cursor position at the end of the primary movement. As with the bias analysis, end-points with a hand angle greater than 3 MADs from the group median were removed to ignore movements that were not target directed. Error ellipsoids were fit per target distance per participant following previous studies (48, 50), giving the angles of the two principal axes and the variability along each axis. The aspect ratio here was calculated as the ratio of standard deviations of the axes aligned more closely to the extent and angle axes respectively. An aspect ratio > 1 indicated movements had more variability in extent than direction. The group average ellipsoid was found by averaging the centre, axis angles and standard deviations over participants.

### Statistical Analysis

Statistical analysis was performed using R (version 4.2.2). In Experiment 1, comparisons were made between contexts by performing Pearson correlations between metrics in the Look and Point context, with paired-samples t-tests used to compare within-subject differences between contexts. Comparisons were also made between all metrics and the asymptotic timeout using Pearson correlations, to understand which might be important predictors of skill on the task. In Experiment 2, improvements in metrics over the experiment was assessed using paired-samples t-tests between the first and last rounds of the experiment. Assessment of the metrics important for overall task performance was done by extracting metrics on the last round of the experiment, after participants had sufficient practice, and entering them into a linear regression where acquire time was predicted by the four kinematic metrics.

A variety of methods were utilised to compare the Look movements to classic findings. Comparisons across distance (movement time scaling, peak speed scaling, and aspect ratio of the error ellipsoids) or angle (bias across target angles) were done using a repeated-measures ANOVA (*afex* package, with Greenhouse-Geisser corrections) and followed up using one-sample or pairwise comparisons on the estimated marginal means (*emmeans* package) with Bonferroni-Holm corrections. The regression of effective index of difficulty on movement time was done using mixed-effect linear regression (*lme4* package), with a fixed effect of effective index of difficulty and random intercepts and slopes per participant, and the p-values of the fixed effects were obtained using Satterthwaite’s approximation (*lmerTest* package). The statistical significance threshold was set at p < .05 throughout.

## Availability

All data and scripts to analyse the data will be made available upon publication of the manuscript.

## Acknowledgements

We thank Jonathan Tsay and Hyosub Kim for helpful comments on the manuscript. Authors F.M and M.M-W were supported by Fellowships from the Alan Turing Institute.

## References

1. H. Tristão, Esports Audience Will Pass Half a Billion in 2022 | Esports Market Analysis. Newzoo (2022) (September 6, 2022).

2. B. Bediou, et al., Meta-analysis of action video game impact on perceptual, attentional, and cognitive skills. Psychol. Bull. 144, 77–110 (2018).

3. C. S. Green, D. Bavelier, Learning, Attentional Control, and Action Video Games. Curr. Biol. 22, R197–R206 (2012).

4. A. Chopin, B. Bediou, D. Bavelier, Altering perception: the case of action video gaming. Curr. Opin. Psychol. 29, 168–173 (2019).

5. S. Green, D. Bavelier, Action-Video-Game Experience Alters the Spatial Resolution of Vision. Psychol. Sci. 18, 88–94 (2007).

6. M. W. G. Dye, C. S. Green, D. Bavelier, Increasing Speed of Processing With Action Video Games. Curr. Dir. Psychol. Sci. 18, 321–326 (2009).

7. G. L. West, N. Al-Aidroos, J. Pratt, Action video game experience affects oculomotor performance. Acta Psychol. (Amst.) 142, 38–42 (2013).

8. C. S. Green, D. Bavelier, Action video game modifies visual selective attention. Nature 423, 534–537 (2003).

9. N. Qiu, et al., Rapid Improvement in Visual Selective Attention Related to Action Video Gaming Experience. Front. Hum. Neurosci. 12, 47 (2018).

10. I. Pedraza-Ramirez, L. Musculus, M. Raab, S. Laborde, Setting the scientific stage for esports psychology: a systematic review. Int. Rev. Sport Exerc. Psychol. 13, 319–352 (2020).

11. M. J. Campbell, A. J. Toth, A. P. Moran, M. Kowal, C. Exton, “eSports: A new window on neurocognitive expertise?” in Progress in Brain Research, (Elsevier, 2018), pp. 161–174.

12. S. Reeves, B. Brown, E. Laurier, Experts at Play: Understanding Skilled Expertise. Games Cult. 4, 205–227 (2009).

13. P. Pickavance, et al., Sensorimotor ability and inhibitory control independently predict attainment in mathematics in children and adolescents. J. Neurophysiol. 127, 1026–1039 (2022).

14. S.-M. Coll, N. E. V. Foster, A. Meilleur, S. M. Brambati, K. L. Hyde, Sensorimotor skills in autism spectrum disorder: A meta-analysis. Res. Autism Spectr. Disord. 76, 101570 (2020).

15. M. C. Rodriguez, et al., Emotional and Behavioral Problems in 4- and 5-Year Old Children With and Without Motor Delays. Front. Pediatr. 7, 474 (2019).

16. A. Mané, E. Donchin, The space fortress game. Acta Psychol. (Amst.) 71, 17–22 (1989).

17. S. Murphy, Video Games, Competition and Exercise: A New Opportunity for Sport Psychologists? Sport Psychol. 23, 487–503 (2009).

18. T. Stafford, M. Dewar, Tracing the Trajectory of Skill Learning With a Very Large Sample of Online Game Players. Psychol. Sci. 25, 511–518 (2014).

19. M. Aung, et al., Predicting Skill Learning in a Large, Longitudinal MOBA Dataset in 2018 IEEE Conference on Computational Intelligence and Games (CIG), (IEEE, 2018), pp. 1–7.

20. J. Huang, E. Yan, G. Cheung, N. Nagappan, T. Zimmermann, Master Maker: Understanding Gaming Skill Through Practice and Habit From Gameplay Behavior. Top. Cogn. Sci. 9, 437–466 (2017).

21. T. Stafford, S. Devlin, R. Sifa, A. Drachen, Exploration and Skill Acquisition in a Major Online Game in The 39th Annual Meeting of the Cognitive Science Society, (2017), p. 7.

22. A. Sapienza, Y. Zeng, A. Bessi, K. Lerman, E. Ferrara, Individual performance in team-based online games. R. Soc. Open Sci. 5, 1–14 (2018).

23. I. Donovan, et al., Assessment of human expertise and movement kinematics in first-person shooter games. Front. Hum. Neurosci. 16 (2022).

24. J. B. Listman, J. S. Tsay, H. E. Kim, W. E. Mackey, D. J. Heeger, Long-Term Motor Learning in the “Wild” With High Volume Video Game Data. Front. Hum. Neurosci. 15, 777779 (2021).

25. A. J. Toth, N. Ramsbottom, C. Constantin, A. Milliet, M. J. Campbell, The effect of expertise, training and neurostimulation on sensory-motor skill in esports. Comput. Hum. Behav. 121, 106782 (2021).

26. A. J. Toth, F. Hojaji, M. J. Campbell, Exploring the mechanisms of target acquisition performance in esports: The role of component kinematic phases on a first person shooter motor skill. Comput. Hum. Behav. 139, 107554 (2023).

27. E. Park, et al., Secrets of Gosu: Understanding Physical Combat Skills of Professional Players in First-Person Shooters in Proceedings of the 2021 CHI Conference on Human Factors in Computing Systems, (ACM, 2021), pp. 1–14.

28. S. K. Coltman, R. J. van Beers, W. P. Medendorp, P. L. Gribble, Sensitivity to error during visuomotor adaptation is similarly modulated by abrupt, gradual, and random perturbation schedules. J. Neurophysiol. 126, 934–945 (2021).

29. O. A. Kim, A. D. Forrence, S. D. McDougle, Learning from the path not taken: Sensory prediction errors are sufficient for implicit adaptation of withheld movements. bioRxiv, 31 (2022).

30. J. Smeets, E. Brenner, Fast corrections of movements with a computer mouse. Spat. Vis. 16, 365–376 (2003).

31. J. S. Tsay, R. B. Ivry, A. Lee, G. Avraham, Moving outside the lab: The viability of conducting sensorimotor learning studies online. Neurons Behav. Data Anal. Theory, 1–22 (2021).

32. I. S. MacKenzie, T. Kauppinen, M. Silfverberg, Accuracy measures for evaluating computer pointing devices in Proceedings of the SIGCHI Conference on Human Factors in Computing Systems – CHI ‘01, (ACM Press, 2001), pp. 9–16.

33. A. K. Mithal, S. A. Douglas, Differences in movement microstructure of the mouse and the finger-controlled isometric joystick in Proceedings of the SIGCHI Conference on Human Factors in Computing Systems Common Ground - CHI ‘96, (ACM Press, 1996), pp. 300–307.

34. G. Phillips, T. J. Triggs, Characteristics of cursor trajectories controlled by the computer mouse. Ergonomics 44, 527–536 (2001).

35. N. Walker, D. E. Meyer, Jo. B. Smelcer, Spatial and temporal characteristics of rapid cursor-positioning movements with electromechanical mice in human-computer interaction. Hum. Factors 35, 431–458 (1993).

36. R. Shadmehr, J. W. Krakauer, A computational neuroanatomy for motor control. Exp. Brain Res. 185, 359–381 (2008).

37. J. Looser, A. Cockburn, J. Savage, On the validity of using First-Person Shooters for Fitts’ law studies. People Comput. XIX 2, 33–36 (2005).

38. D. Elliott, et al., Goal-directed aiming: Two components but multiple processes. Psychol. Bull. 136, 1023–1044 (2010).

39. W. Abend, E. Bizzi, P. Morasso, Human arm trajectory formation. Brain J. Neurol. 105, 331–348 (1982).

40. P. Morasso, Spatial control of arm movements. Exp. Brain Res. 42 (1981).

41. C. Atkeson, J. Hollerbach, Kinematic features of unrestrained vertical arm movements. J. Neurosci. 5, 2318–2330 (1985).

42. L. E. Sergio, S. H. Scott, Hand and joint paths during reaching movements with and without vision. Exp. Brain Res. 122, 157–164 (1998).

43. A. Kistemaker, J. D. Wong, P. L. Gribble, The cost of moving optimally: kinematic path selection. J. Neurophysiol. 112, 1815–1824 (2014).

44. J. S. Brown, A. T. Slater-Hammel, Discrete movements in the horizontal plane as a function of their length and direction. J. Exp. Psychol. 39, 84 (1949).

45. P. M. Fitts, The information capacity of the human motor system in controlling the amplitude of movement. J. Exp. Psychol. 47, 381 (1954).

46. R. W. Soukoreff, I. S. MacKenzie, Towards a standard for pointing device evaluation, perspectives on 27 years of Fitts’ law research in HCI. Int. J. Hum.-Comput. Stud. 61, 751–789 (2004).

47. I. S. MacKenzie, C. Ware, Lag as a determinant of human performance in interactive systems in Proceedings of the SIGCHI Conference on Human Factors in Computing Systems - CHI ‘93, (ACM Press, 1993), pp. 488–493.

48. J. Gordon, M. F. Ghilardi, C. Ghez, Accuracy of planar reaching movements: I. Independence of direction and extent variability. Exp. Brain Res. 99, 97–111 (1994).

49. C. M. Harris, D. M. Wolpert, Signal-dependent noise determines motor planning. Nature 394, 780–784 (1998).

50. J. Messier, J. F. Kalaska, Comparison of variability of initial kinematics and endpoints of reaching movements. Exp. Brain Res. 125, 139–152 (1999).

51. G. H. Begbie, Accuracy of Aiming in Linear Hand-Movements. Q. J. Exp. Psychol. 11, 65–75 (1959).

52. F. Ghilardi, J. Gordon, C. Ghez, Learning a visuomotor transformation in a local area of work space produces directional biases in other areas. J. Neurophysiol. 73, 2535–2539 (1995).

53. A. Chandy, et al., Motor Biases are Persistent and Consistent (2021).

54. J. J. Thompson, M. R. Blair, L. Chen, A. J. Henrey, Video Game Telemetry as a Critical Tool in the Study of Complex Skill Learning. PLoS ONE 8, e75129 (2013).

55. J. J. Thompson, C. M. McColeman, E. R. Stepanova, M. R. Blair, Using Video Game Telemetry Data to Research Motor Chunking, Action Latencies, and Complex Cognitive-Motor Skill Learning. Top. Cogn. Sci. 9, 467–484 (2017).

56. J. A. Adams, Warm-up Decrement in Performance on the Pursuit-Rotor. Am. J. Psychol. 65, 404 (1952).

57. A. Smith, A. Ghazizadeh, R. Shadmehr, Interacting Adaptive Processes with Different Timescales Underlie Short-Term Motor Learning. PLoS Biol. 4, e179 (2006).

58. G. S. Snoddy, Learning and stability. J. Appl. Psychol. 10, 1–36 (1926).

59. A. Fishbach, S. A. Roy, C. Bastianen, L. E. Miller, J. C. Houk, Deciding when and how to correct a movement: discrete submovements as a decision making process. Exp. Brain Res. 177, 45–63 (2007).

60. D. Lee, N. L. Port, A. P. Georgopoulos, Manual interception of moving targets II. On-line control of overlapping submovements: II. On-line control of overlapping submovements. Exp. Brain Res. 116, 421–433 (1997).

61. T. E. Milner, M. M. Ijaz, The effect of accuracy constraints on three-dimensional movement kinematics. Neuroscience 35, 365–374 (1990).

62. J. Pratt, A. L. Chasteen, R. A. Abrams, Rapid aimed limb movements: age differences and practice effects in component submovements. Psychol. Aging 9, 325 (1994).

63. R. A. Abrams, J. Pratt, Rapid Aimed Limb Movements: Differential Effects of Practice on Component Submovements. J. Mot. Behav. 25, 288–298 (1993).

64. D. E. Meyer, R. A. Abrams, S. Kornblum, C. E. Wright, J. Keith Smith, Optimality in human motor performance: ideal control of rapid aimed movements. Psychol. Rev. 95, 340 (1988).

65. S. Zhai, Characterizing computer input with Fitts’ law parameters—the information and non-information aspects of pointing. Int. J. Hum.-Comput. Stud. 61, 791–809 (2004).

66. D. McNamee, D. M. Wolpert, Internal Models in Biological Control. Annu. Rev. Control Robot. Auton. Syst. 2, 339–364 (2019).

67. S. H. Scott, The computational and neural basis of voluntary motor control and planning. Trends Cogn. Sci. 16, 541–549 (2012).

68. J. B. Heald, M. Lengyel, D. M. Wolpert, Contextual inference underlies the learning of sensorimotor repertoires. Nature 600, 489–493 (2021).

69. D. M. Wolpert, M. Kawato, Multiple paired forward and inverse models for motor control. Neural Netw. 11, 1317–1329 (1998).

70. T. P. Lillicrap, et al., Adapting to inversion of the visual field: a new twist on an old problem. Exp. Brain Res. 228, 327–339 (2013).

71. S. Telgen, D. Parvin, J. Diedrichsen, Mirror Reversal and Visual Rotation Are Learned and Consolidated via Separate Mechanisms: Recalibrating or Learning De Novo? J. Neurosci. 34, 13768–13779 (2014).

72. S. A. Wilterson, J. A. Taylor, Implicit Visuomotor Adaptation Remains Limited after Several Days of Training. eneuro 8, ENEURO.0312-20.2021 (2021).

73. S. Yang, N. J. Cowan, A. M. Haith, De novo learning versus adaptation of continuous control in a manual tracking task. eLife 10, e62578 (2021).

74. M. Desmurget, S. Grafton, Forward modeling allows feedback control for fast reaching movements. Trends Cogn. Sci. 4, 423–431 (2000).

75. R. C. Miall, D. M. Wolpert, Forward Models for Physiological Motor Control. Neural Netw. 9, 1265–1279 (1996).

76. A. Buneo, M. R. Jarvis, A. P. Batista, R. A. Andersen, Direct visuomotor transformations for reaching. Nature 416, 632–636 (2002).

77. M. Desmurget, et al., Role of the posterior parietal cortex in updating reaching movements to a visual target. Nat. Neurosci. 2, 563–567 (1999).

78. J. W. Krakauer, Z. M. Pine, M.-F. Ghilardi, C. Ghez, Learning of Visuomotor Transformations for Vectorial Planning of Reaching Trajectories. J. Neurosci. 20, 8916–8924 (2000).

79. J. Wang, R. L. Sainburg, Adaptation to Visuomotor Rotations Remaps Movement Vectors, Not Final Positions. J. Neurosci. 25, 4024–4030 (2005).

80. H. G. Wu, M. A. Smith, The Generalization of Visuomotor Learning to Untrained Movements and Movement Sequences Based on Movement Vector and Goal Location Remapping. J. Neurosci. 33, 10772–10789 (2013).

81. H. Tanaka, T. J. Sejnowski, J. W. Krakauer, Adaptation to Visuomotor Rotation Through Interaction Between Posterior Parietal and Motor Cortical Areas. J. Neurophysiol. 102, 2921–2932 (2009).

82. S. F. W. Neggers, H. Bekkering, Ocular Gaze is Anchored to the Target of an Ongoing Pointing Movement. J. Neurophysiol. 83, 639–651 (2000).

83. C. Prablanc, J. F. Echallier, E. Komilis, M. Jeannerod, Optimal response of eye and hand motor systems in pointing at a visual target. Biol. Cybern. 35, 113–124 (1979).

84. A. Kenny, H. Koesling, D. Delaney, S. McLoone, T. Ward, A Preliminary Investigation into Eye Gaze Data in a First Person Shooter Game in Proceedings of the 19th European Conference on Modelling and Simulation, (2005), p. 6.

85. H. Alayed, F. Frangoudes, C. Neuman, Behavioral-based cheating detection in online first person shooters using machine learning techniques in 2013 IEEE Conference on Computational Inteligence in Games (CIG), (IEEE, 2013), pp. 1–8.

86. L. Galli, D. Loiacono, L. Cardamone, P. L. Lanzi, A cheating detection framework for Unreal Tournament III: A machine learning approach in 2011 IEEE Conference on Computational Intelligence and Games (CIG’11), (IEEE, 2011), pp. 266–272.

87. H.-K. Pao, K.-T. Chen, H.-C. Chang, Game Bot Detection via Avatar Trajectory Analysis. IEEE Trans. Comput. Intell. AI Games 2, 162–175 (2010).

88. K. Waananen, Touring Team Liquid and Alienware’s The Pro Lab. Esports Insid. (2022) (November 21, 2022).

89. J. Brookes, M. Warburton, M. Alghadier, M. Mon-Williams, F. Mushtaq, Studying human behavior with virtual reality: The Unity Experiment Framework. Behav. Res. Methods 52, 455–463 (2019).

